# Feedback between membrane tension, lipid shape and curvature in the formation of packing defects

**DOI:** 10.1101/389627

**Authors:** M. Pinot, S. Vanni, E. Ambroggio, D. Guet, B. Goud, J.-B. Manneville

## Abstract

Lipid packing defects favor the binding of proteins to cellular membranes by creating spaces between lipid head groups that allow the insertion of amphipathic helices or lipid modifications. The density of packing defects in a lipid membrane is well known to increase with membrane curvature and in the presence of conical-shaped lipids. In contrast, the role of membrane tension in the formation of lipid packing defects has been poorly investigated. Here we use a combination of numerical simulations and experiments to measure the effect of membrane tension on the density of lipid packing defects. We first monitor the binding of ALPS (amphipathic lipid packing sensor) to giant unilamellar vesicles and observe a striking periodic binding of ALPS that we attribute to osmotically-induced membrane tension and transient membrane pore formation. Using micropipette aspiration experiments, we show that a high membrane tension induces a reversible increase in the density of lipid packing defects. We next focus on packing defects induced by lipid shape and show that conical lipids generate packing defects similar to that induced by membrane tension and enhance membrane deformation due to the insertion of the ALPS helix. Both cyclic ALPS binding and the cooperative effect of ALPS binding and conical lipids on membrane deformation result from an interplay between helix insertion and lipid packing defects created by membrane tension, conical lipids and/or membrane curvature. We propose that feedback mechanisms involving membrane tension, lipid shape and membrane curvature play a crucial role in membrane deformation and intracellular transport events.

## Introduction

Feedback loops are fundamental features of many diverse cellular functions, from pattern formation (1–3) and establishment of cell shape, polarity and migration (4–6) to circadian rhythms (7) and the cell cycle (8) or membrane trafficking (9). Mechanisms based on reaction-diffusion and activation/inhibition processes generate positive and negative feedback loops that lead either to a stable steady-state or to oscillating regimes.

In the case of membrane trafficking, membrane deformation and fission, the first steps in the formation of a transport carrier (10), could be driven by feedback mechanisms. The macroscopic physical characteristics of a membrane - curvature, bending rigidity and tension - couples to microscopic parameters such as the lipid composition and packing (11, 12), and positive feedback loops could amplify an initially small membrane deformation to generate budding and ultimately lead to scission. For instance, feedback between actin and curvature-sensitive endocytic proteins such as dynamin or BAR (Bin/Amphiphysin/Rvs167) domain-containing proteins has been proposed to participate in budding and scission during endocytosis (13–15). Similarly, during cell migration, membrane tension was shown to impact on actin dynamics (16) and to couple to membrane bending via the F-BAR domain protein FBP17 (17). Intracellular coat proteins have also been shown to induce budding and prepare membranes for fission by feedback-based mechanisms (18, 19).

Central to this coupling between membrane physical parameters is the notion of lipid packing. Lipid packing is defined by the density of lipid molecules within the bilayer. Regions where this density decreases are called ‘lipid packing defects’ (20, 21). Lipid packing defects can appear due to membrane bending or, in a flat membrane, due to the presence of conical lipids (22–24). Stretching the membrane should in principle also induce the formation of lipid packing defects (11) although convincing evidence is still lacking. Here we study the roles of membrane curvature, lipid shape and membrane tension in the generation of lipid packing defects in the context of membrane budding and fission. We use the Amphipathic Lipid Packing Sensor (ALPS) motifs from the COPI (coat protein complex I) coat-associated protein ArfGAP1 (Arf GTPase activating protein 1) to detect lipid packing defects (25), and to simultaneously test for potential membrane bending due to shallow hydrophobic helix insertion. Based on our results, we propose that feedback loops play a critical role in membrane deformation. We suggest 1) that a high membrane tension can feedback on amphipathic helix insertion by increasing the density of lipid packing defects; and 2) that lipid packing defects generated by conical lipids and amplified by curvature could lead to instability in curved membranes and ultimately to membrane scission.

## Materials and methods

### Reagents

Egg phosphatidylcholine (EPC), liver phosphatidylinositol (LPI), liver phosphatidylethanolamine (LPE), brain phosphatidylserine (BPS), cholesterol, 1,2- dioleoyl-*sn*-glycero-3-phosphocholine (DOPC), brain sphingomyelin (BSM), brain polar lipid (BPL), 1-stearoyl-2-docosahexaenoyl-sn-glycero-3-phosphoethanolamine and 1-2- dioleoyl-sn-glycerol (DAG) were from Avanti Polar Lipids. The fluorescent lipid DHPE-TexasRed and the Alexa-maleimide fluorophores were from Invitrogen.

### Peptide and protein purification

Purification of the ArfGAP1 lipid packing sensing tandem motif ALPS1-ALPS2 (here denoted as ALPS) and of ArfGAP1 and fluorescent labeling of ALPS with the Alexa 488 or Alexa 546 dyes were performed using the protocol described previously (19) (see Supplementary Methods).

### Preparation of giant unilamellar vesicles

GUVs were grown at room temperature on ITO slides using the electroformation technique described previously (26). To avoid any osmotic shock, the osmotic pressure of the sucrose solution was adjusted to match that of the buffers containing the proteins, HKM buffer (50mM Hepes pH7.2, 120 mM Potassium Acetate, 1 mM MgCl_2_) for ALPS experiments and HKM supplemented with 2 mM EDTA for ArfGAP1 experiments. GUVs were grown in 280 mOsm sucrose for ALPS experiments and in 320 mOsm sucrose for ArfGAP1 experiments. For hypertonic and hypotonic experiments with ALPS, GUVs were grown in 260 mOsm sucrose (corresponding to an osmotic pressure difference ΔΠ = +20 mOsm) and 320 mOsm sucrose (corresponding to an osmotic pressure difference ΔΠ = −40 mOsm) respectively. GUVs containing diacylglycerols were grown from a ‘Golgi mix’ composed of 50% (mol/mol) EPC, 19% LPE, 5% BPS, 10% LPI, and 16% cholesterol and supplemented with 0.1% to 20% (mol/mol) DAG. GUVs containing PUFAs (polyunsaturated fatty acid) were grown from a ‘DAG-Golgi mix’ supplemented with 1% to 10% (mol/mol) 1-stearoyl-2-docosahexaenoyl-sn-glycero-3-phosphoethanolamine. GUV membranes were made fluorescent by adding 1% (vol/vol) DHPE-TexasRed. For micropipette aspiration, GUVs were grown from DOPC without fluorescent phospholipid.

### Micropipet aspiration experiments

A 100–200 μl open micromanipulation chamber was built with two clean glass coverslips as described in (27). The surfaces of the chamber and of the micropipette were incubated with 10 mg/ml β-casein for 10 min and then rinsed with HKM buffer to prevent adhesion of the GUVs to the glass. 100–200 μl of 1 μM ALPS-Alexa 488 in HKM buffer were injected into the chamber to monitor ALPS binding. 5 μl of GUVs were transferred from the growth chamber to the micromanipulation chamber. The chamber was sealed with mineral oil to prevent water evaporation. Membrane tension σ was determined from σ = (Δ*P* • *R_pip_*)/2(1 – *R_pip_/R_ves_*) where *R_pip_* is the pipette radius, *R_ves_* is the vesicle radius, and Δ*P* is the difference of hydrostatic pressure caused by the vertical displacement of the water reservoir linked to the pipette. Membrane tension of the GUV varied from 5.10^-6^ to 3.10^-4^ N/m (28). Images were taken on a Nikon A1R Ti-Eclipse inverted confocal microscope or on a Zeiss LSM510 Meta confocal microscope.

### Molecular dynamics (MD) simulations

MD simulations were performed using the coarse-grain (CG) lipid model by Klein and coworkers (29–31). All simulations were performed with the LAMMPS software (32). Configurations for the various systems were generated by converting atomistic snapshots using the CG-it software (https://github.com/CG-it). Pressure and temperature were controlled using a Nosé-Hoover thermostat (33) and barostat (34, 35), with target temperature and pressure of 300 K and 1 atm, respectively. For surface tension simulations, the lateral *xy* dimensions of the systems were constrained to be equal and fixed, while the orthogonal dimension *z* was allowed to fluctuate independently. Van der Waals and electrostatics were truncated at 1.5 nm, with long-range electrostatics beyond this cutoff computed using the particle-particle-particle-mesh (PPPM) solver with an RMS force error of 10^-5^ kcal mol^-1^ Å^-1^ and order 3 (36). In all simulations, a time step of 20 fs was used. The DOPC bilayer system consisted of 1296 DOPC molecules per leaflet (2592 DOPC molecules in total), and each system was run for 400 ns. Analyses were performed on the last 300 ns of each MD run. All molecular graphics were generated with VMD (37). Density profiles were computed by averaging the mass density of the phosphate groups along the *z* axis using a grid of 0.2 nm resolution.

For all simulations, surface tension was computed from the diagonal values of the pressure tensor (*P_xx_, P_yy_* and *P_zz_*) using the Kirkwood-Irving method (38):

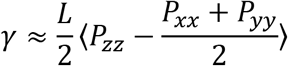

where *L* is the box length in the *z* dimension and 〈…〉 means an ensemble average.

Lipid-packing defects were computed using a previously described algorithm (22, 25), that is described in detail in http://packmem.ipmc.cnrs.fr. In brief, the membrane is mapped to a grid (with a granularity of 1 Å) parallel to the plane of the bilayer and each grid point is scanned vertically starting from outside of the membrane towards its interior. If only aliphatic atoms are encountered, the grid point is considered as an elementary packing defect of 1 Å^2^. Only deep defects (below 2 Å of the glycerol level) were considered in this study. Adjacent elementary defects are then clustered using a connected component algorithm, and the area of each cluster is calculated (a packing defect is thus a cluster of elementary defects). This analysis is done for each leaflet separately. Statistics are then accumulated over all frames of a trajectory following the same procedure. The obtained distributions are then fit to a mono-exponential decay: *p*(*A*) = *be*^-*A/π*_*type*_^, where *p*(*A*) is the probability of finding a defect of area *A* Å^2^, *π_type_* is the packing defect constant in units of Å^2^ that is reported in Fig. 2E, and *b* is a constant. The fit is done on defects larger than 15 Å^2^ and for probabilities larger or equal to 10^-4^. The higher the packing-defect constant, the higher the probability of finding large defects.

### Fluorescence quantification

The amount of fluorescently labeled ALPS bound to the membrane of the GUVs was quantified using the OvalProfile plug-in in ImageJ (http://rsb.info.nih.gov/ij). Briefly, the maximum intensity value along a radius starting from the center of the GUV was measured. This measurement was repeated every degree to generate the circumferential intensity profile along the GUV membrane. ALPS binding was defined as the averaged value of the circumferential intensity profile to which background ALPS fluorescence was subtracted.

### Measurement of sorting coefficients

Analyses of confocal fluorescence images were performed using a protocol described previously (39). For each GUV, the membrane contour was transformed into a line using a home-made Matlab code. The mean fluorescence intensities of lipid and ALPS labelling in the GUV was estimated using the width at half the maximum of the membrane profile, *w*, by averaging the intensity for pixels in the 1*w*-range. The mean background level of each fluorescence channel was estimated from pixels localized outside of the GUV at a distance between 2*w* and 6*w*. The fluorescence of membrane deformations and invaginations was measured by manually selecting all pixels in deformations and invaginations where lipid signal was present outside of the GUV. The mean intensities of the two channels (ALPS and lipid labelling) were measured on GUV and membrane invaginations and the sorting coefficient was calculated as *Sorting* = (*I_ALPS_/I_hipid_*)_tube_/(*I_ALPS_/I_lipid_*)_*GUV*_.

## Results

### Heterogeneous and time-dependent ALPS binding to GUV membranes suggests a role for membrane tension in lipid packing defect formation

We first measured the binding of fluorescently labeled ALPS, a quantitative indicator of the density of lipid packing defects, to giant unilamellar vesicles (GUVs) made of pure 1,2- dioleoyl-sn-glycero-3-phosphocholine (DOPC). DOPC should favour ALPS binding due to its weakly conical shape of DOPC (40). Surprisingly, we found that the binding of ALPS was very heterogeneous among the GUV population. Within the same confocal field, we could observe GUVs exhibiting a strong fluorescent ALPS signal while no signal could be detected from others (Fig. S1A in the Supporting Material). The intensity of ALPS binding did not correlate with the size of the GUVs. Even more surprisingly, some GUVs displayed a time-dependent fluorescent signal (Movie S1 and Fig. S1 in the Supporting Material), suggesting that the density of lipid packing defects varied with time. In about 10% of the GUVs, the fluorescent signal was periodic (Fig. 1A and S1A). The fluorescence increased with time up to a maximal level then abruptly dropped before increasing linearly again (Fig. 1B). The period of the oscillations was ≈ 100-500 s. We could observe 2-3 oscillations during the course of the time-lapse acquisition. Since neither membrane composition nor membrane curvature significantly changed during the experiment, membrane tension could be responsible for these variations in lipid packing defect density. In the Canham-Helfrich-Evans elastic theory of membranes, a low membrane tension unfolds the excess area stored in membrane shape fluctuations while higher tensions stretch the membrane (28). Stretching at high tension could thus increase the spacing between lipid head groups and thus induce the formation of lipid packing defects and increase ALPS binding. To explain the oscillatory binding of ALPS, we hypothesized that an osmotically-driven increase in membrane tension leads to the formation of transient pores that release the tension (41–43). Accordingly, the GUV diameter followed the same oscillating pattern as the fluorescence signal (Fig. 1C). The drop in fluorescence coincided with a drop in GUV radius. The maximal increase in GUV diameter was of a few tenths of micrometres corresponding to about 3-4% of the typical GUV diameter. To test our hypothesis we monitored ALPS binding in hyperosmotic or hypoosmotic buffers. Consistent with our hypothesis, in a hypoosmotic buffer which increases the initial GUV tension, the period of the oscillations decreased (Fig. 1D and S1B). We also performed experiments in the presence of a soluble fluorophore (fluorescein or sulforhodamin) to assess the formation of membrane pores. The fluorescent signal, initially present only in the external buffer, progressively entered inside GUVs in the presence of ALPS with a step increase after each drop in ALPS binding (Fig. 1E and Movie 2 in the Supporting Material), suggesting that transient pores open in the GUV membrane and/or that the bilayer permeability increases.

**Figure 1.**
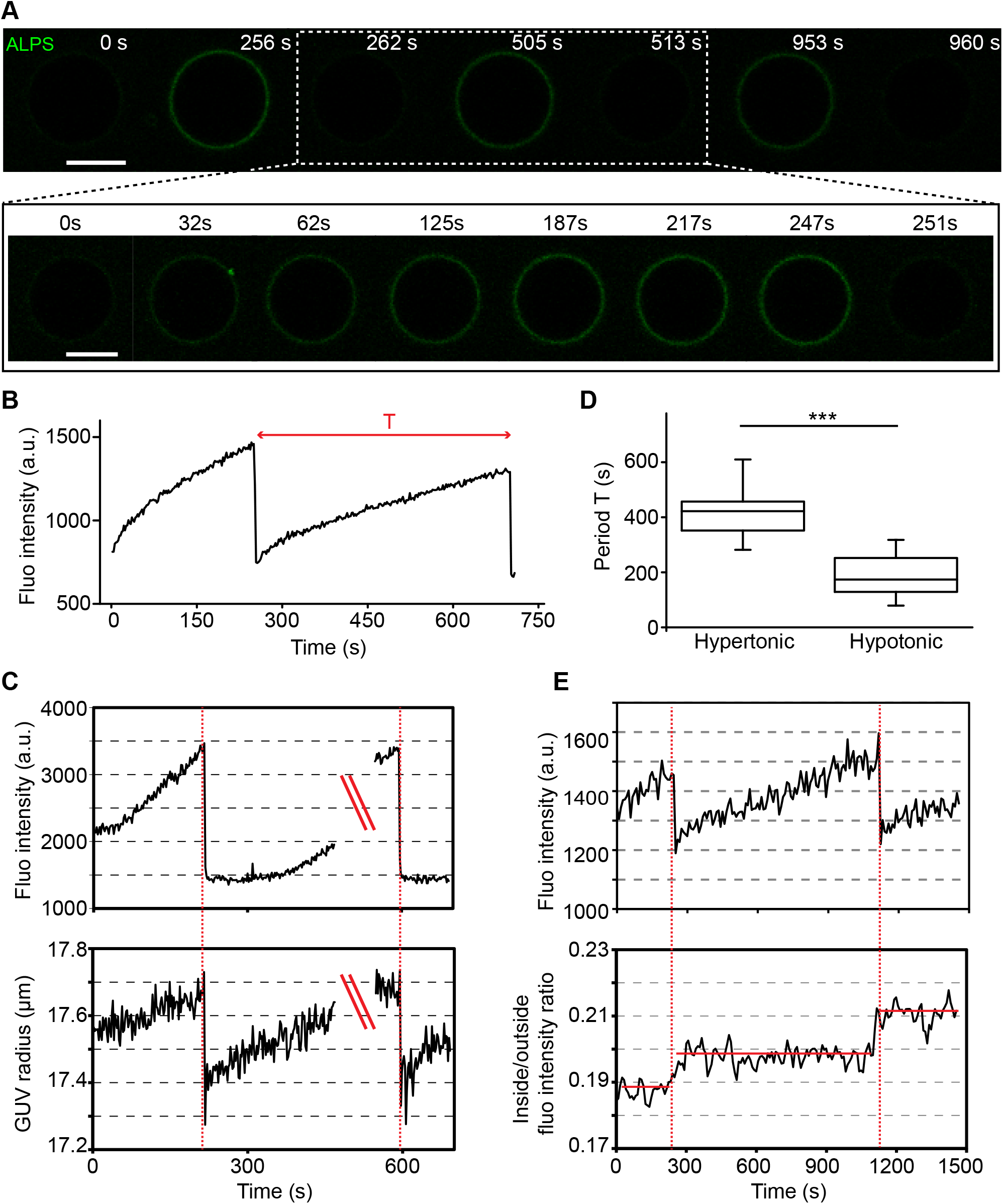
Periodic ALPS binding to GUVs membranes. (A) Spontaneous oscillations of ALPS binding (1 μM) on GUVs made of pure DOPC. ALPS is shown in green. Time series of ALPS binding. See also Movie 1 in the Supporting Material. (B) Measurement of ALPS binding as a function of time. Measurements shown were performed on the data shown in A. (C) Correlation between ALPS binding and GUV diameter. A drop in ALPS binding corresponds to a drop in GUV diameter. The double red bar indicates that measurements were interrupted. (D) Quantification of the period of the oscillations in hypertonic (osmotic pressure difference ΔΠ = *+20* mOsm) or hypotonic conditions (ΔΠ = *-40* mOsm). Data shown are averages of *N=20* GUVs from four independent experiments. The error bars represent standard deviations and the box the 70 *%* probability. Data were analyzed using a Student’s t test; *** indicate P<0.0001. (E) Correlation between ALPS binding and entry of a soluble fluorophore (fluorescein or sulforhodamin) inside the GUV. A drop in ALPS binding corresponds to an increase in the fluorescent dye signal inside the GUV, suggesting that transient pores successively open and close in the membrane. See also Movie 2 in the Supporting Material. Scale bars, 10 μm.

### ALPS binding is modulated by membrane tension

To further test a potential role of membrane tension in the generation of lipid packing defects, we used micropipette aspiration of GUVs to control membrane tension. ALPS binding mirrored the variations in tension imposed on the GUV membrane by the applied pressure difference between the inside and the outside of the pipet. The fluorescent changes were reversible since when the tension was released, the fluorescence decreased to levels similar to those observed prior to aspiration. We could perform cycles of binding and unbinding of ALPS by alternating the tension between ~0.1 mN/m and ~0.3 mN/m (Fig. 2A) similar to the spontaneous oscillatory binding observed above. These results demonstrate a direct role for membrane tension in modulating the binding of ALPS to lipid membranes.

**Figure 2.**
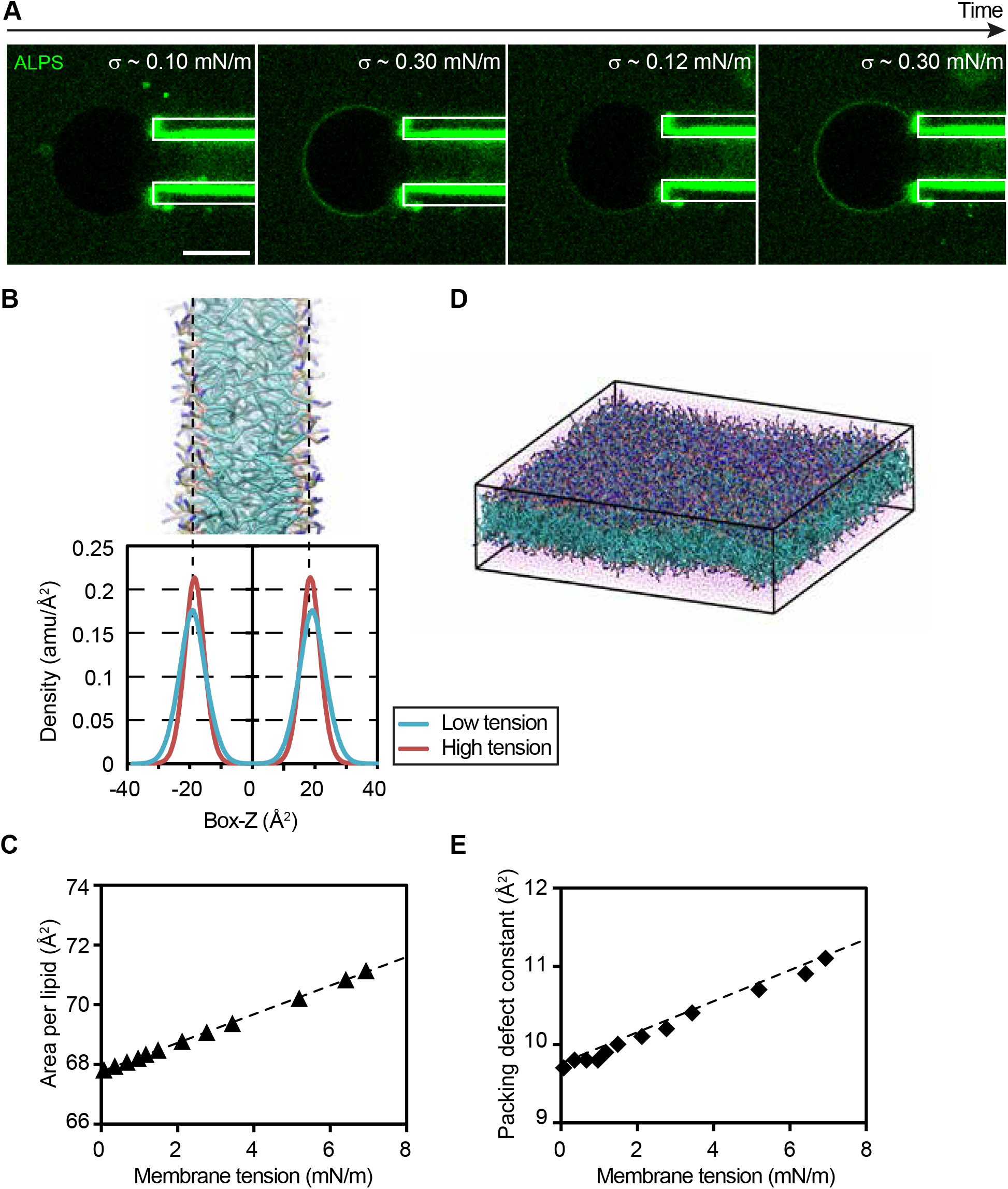
The density of lipid packing defects is modulated by membrane tension. (A) Typical micropipette aspiration experiment showing how the binding of ALPS (1 μM) correlates with variations in membrane tension. Scale bars, 10 μm. (B-E) Coarse grained molecular dynamics simulations of the effect of membrane tension on lipid packing defects. (B) Lateral view of a DOPC lipid bilayer and corresponding density profile for the phosphate group at low and high membrane tension. (C) Increase in area per lipid as a function of membrane tension. The relationship between these two parameters is linear in the range of membrane tensions investigated (D). Representative snapshot of the DOPC lipid bilayers used for all the analyses. (E) Relationship between surface tension and lipid packing defects in the investigated DOPC bilayers. The density of lipid packing defects increases with surface tension. In all plots, error bars (not shown) are below 0.1 Å^2^. Color code: water, red; acyl chains, cyan; glycerol, pink, phosphate group, brown, choline group, blue.

### Membrane tension increases the density of lipid packing defects

ALPS binding to DOPC GUVs increases at elevated membrane tension (Figs. 1 and 2A). This observation suggests that lateral membrane tension increases the density of lipid packing defects. To investigate the molecular consequences of variations in surface tension on the abundance of lipid-packing defects, we performed coarse-grained molecular dynamics (MD) simulations on DOPC lipid bilayers.

As expected from the Canham-Helfrich-Evans elastic theory, increasing surface tension decreases membrane undulations, as can be observed from the lateral density profile of the phosphate groups of the DOPC molecules in the bilayer (Fig 2B). This effect is coupled with a linear increase in area per lipid for increasing surface tensions (Fig. 2C). Hence, to avoid possible finite-size effects resulting from the artificial suppression of bilayer vibrational modes in the MD simulations (44), we performed our MD on a large system of approximately 30 nm x 30 nm in size, and consisting of 2592 DOPC molecules (Fig. 2D).

We next computed lipid-packing defects as a function of surface tension using a previously described Cartesian algorithm (22, 23, 25). As can be seen in Fig. 2E, lipid-packing defects appear to increase linearly as a function of surface tension. Since (i) ALPS binding has been shown to take place above a specific threshold (23) and (ii) bilayers consisting of DOPC lipids are a likely substrate for the binding of ALPS motifs, our MD simulations suggest that even a small increase in membrane tension might be sufficient to trigger ALPS binding to DOPC membranes.

### Conical lipids and membrane tension increase ALPS binding in a similar fashion

The role of membrane tension in the generation of lipid packing defects has only been poorly investigated to date. We showed in figures 1 and 2 that a high membrane tension facilitates the formation of lipid packing defects. The density of packing defects in a lipid bilayer is also known to depend on membrane curvature and on lipid geometrical shape (22, 25). In the following, we first compare how lipid shape and membrane tension influence lipid packing and then investigate how feedbacks between lipid shape and membrane curvature could induce membrane deformation. As expected, the presence of conical lipids (the diacylglycerol di-oleyl-glycerol DOG) increased the density of lipid packing defects as shown by an increased binding of ALPS on GUVs containing DOG (Fig. S2A-C in the Supporting Material). The levels of ALPS binding were comparable to those reached when a high membrane tension was imposed on the GUV by micropipette aspiration (Fig. S2D-F), showing that conical lipids and membrane tension can generate lipid packing defects to similar extents.

### Conical lipids facilitate ALPS binding and subsequent membrane deformation

To get more insight into the interplay between lipid shape and packing defects, we quantified ALPS binding in the presence of increasing amounts of DOG (Fig. 3A-C). ALPS binding increased with the proportion of conical lipids for all concentrations of ALPS we tested within the 1 nM – 3 μM range (Fig. 3C). We next added polyunsaturated fatty acids (PUFAs) to compete with lipid packing defects induced by DOG. PUFAs are known to reduce the density of lipid packing defects by adapting their conformation to local lipid packing (45, 46). The addition of 2% PUFAs to 5% DOG-containing GUVs induced a twofold decrease in ALPS binding while adding 10% PUFA almost completely abolished ALPS binding (Fig. S3A-B in the Supporting Material). We further confirmed the role of lipid shape in the generation of lipid packing defects by producing *in situ* conical lipids from GUVs made of DOPC using phospholipases (Fig. S3C). Production of DOPA by Phospholipase D (PLD) or production of DOG by Phospholipase C (PLC) both induced a time-dependent increase in ALPS binding, with the more conical lipid DOG inducing a faster increase in ALPS binding. DOG production by PLC ultimately led to GUV membrane destabilization, rupture and fission (Fig. S3C and Movie S3 in the Supporting Material), as reported previously (47).

**Figure 3.**
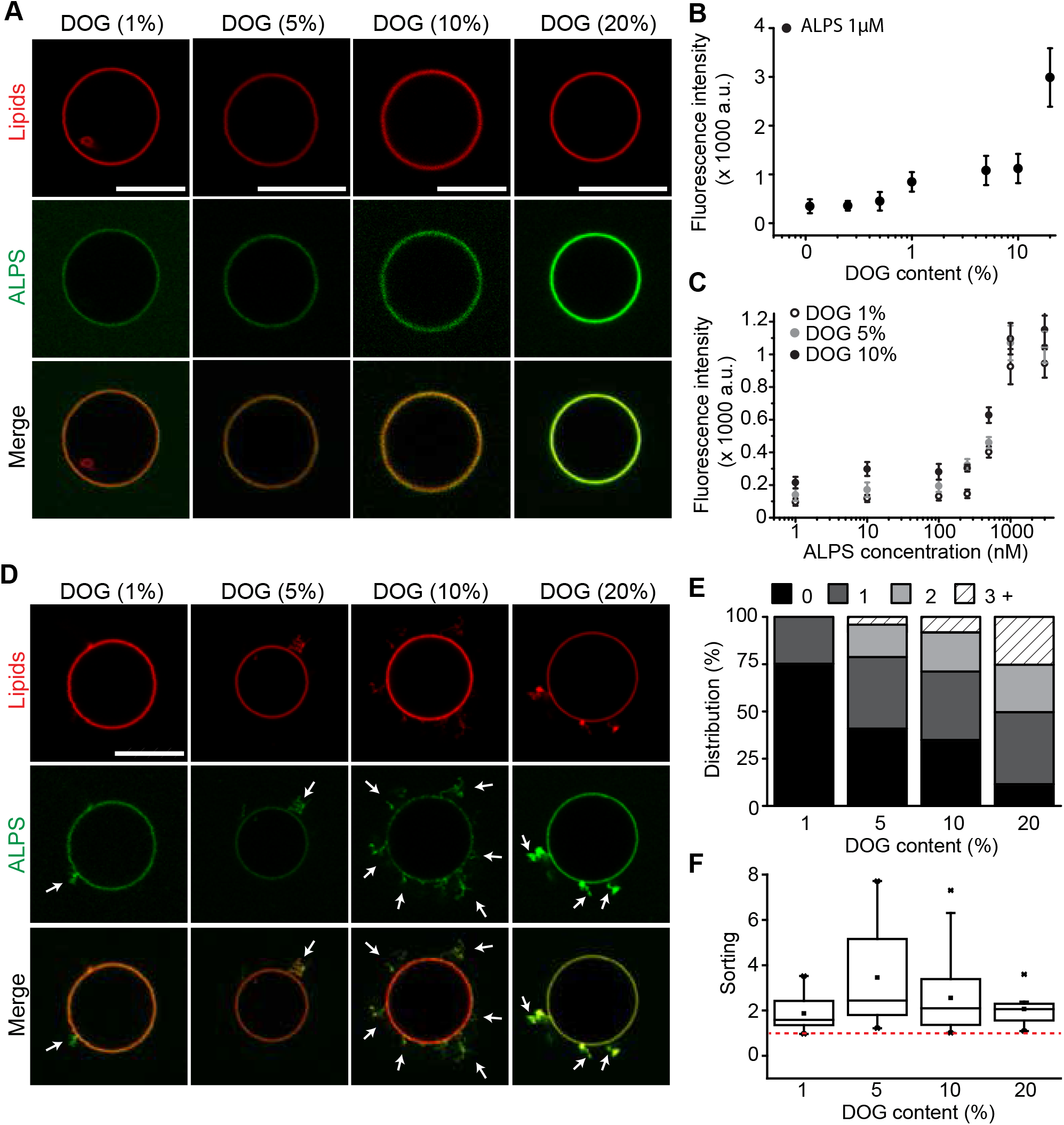
Conical lipids facilitate the formation of packing defects and membrane deformation induced by ALPS binding. (A) ALPS binding (1 μM, green) to GUVs increases with increasing amounts of DOG. Note that the lipid membrane fluorescence (red) may vary depending on the efficiency of the incorporation of the DHPE-TexasRed lipid dye. (B) Quantification of ALPS binding as a function of DOG content. Fluorescence of membrane bound ALPS was measured relative to the background ALPS fluorescence outside the GUV. (C) ALPS binding to GUVs as a function of ALPS bulk concentration. The curves display two regimes: at bulk concentrations of ALPS below 0.3-0.5 μM, ALPS binding is low and no membrane deformation is observed while at concentrations above 0.3-0.5 μM, ALPS binding is high and membrane deformation is observed. Note the logarithmic scale on the x-axis in (B) and (C). (D) Membrane deformation induced by ALPS binding (1 μM) increases with increasing amounts of DOG. ALPS fluorescence is shown in green and the GUV membrane in red. (E) Quantification of membrane deformation as a function of DOG content. GUVs were classified in four categories: GUVs showing no visible deformation and GUVs showing 1, 2 or three and more deformations profiles. Data were pooled from N=381 deformations in 65 GUVs from 8 independent experiments. (F) Sorting of ALPS in the curved membrane deformations. The quantification shows the sorting coefficient as defined by the ratio of the fluorescence of ALPS (green channel) normalized by the lipid membrane fluorescence (red channel) measured in the deformations divided by the same quantity measured in the flat GUV membrane (see Materials and Methods). A sorting value above 1 indicates enrichment of ALPS in the membrane deformations. The box plot shows the 25-75^th^ percentile (box), the 1-99^th^ percentile (bars), the median value (horizontal line), the mean value average (square) and the maximal or minimal (stars) values of the distribution. Scale bars, 10 μm.

As for other membrane insertions, ALPS binding to membranes is expected to induce membrane curvature and tubulation in accordance to the bilayer-couple hypothesis (10, 48). ALPS binding displayed a typical sigmoid curve as a function of ALPS concentration (Fig. 3C), suggesting that two binding regimes exist, a low binding regime below 0.5 μM in which ALPS does not induce membrane deformation and a high binding regime above 0.5 μM in which ALPS induces membrane deformation, as reported for other amphipathic helices (49). Indeed at concentrations above 1μM, both ALPS and ArfGAP1 were able to form tubules on flat membrane sheets or small liposomes (Fig. S4A, B and Movies S4 and S5 in the Supporting Material). Membrane deformation by 1 μM ALPS was enhanced in the presence of increasing amounts of DOG (Fig. 3D-E). With 10% to 20% DOG, extensive membrane deformation could be observed in the presence of ALPS or ArfGAP1 (Fig. S4C, D). Interestingly, ALPS was significantly enriched in the membrane deformations compared to the flat GUV membrane at all tested DOG concentrations (Fig. 3F), consistent with our previous finding that ALPS binding sharply increases on curved lipid nanotubes (19).

## Discussion

We have used a combination of *in vitro* experiments on model membranes and numerical simulations to study the interplay between membrane tension, lipid shape and lipid packing. We have used the amphipathic lipid packing sensor (ALPS) tandem motif of ArfGAP1 as a reporter of lipid packing defect density. ALPS was shown previously to bind preferentially to membranes exhibiting a high density of lipid packing defects induced either by a high membrane curvature (19, 23, 50) or the presence of conical lipids (22, 23, 25, 50). Such preferential binding is driven by changes in intra-membrane stresses impacting the lateral pressure profile (10, 51). Here we focus on the effects of membrane tension and lipid shape on potential changes in intra-membrane stresses. We show that a high membrane tension facilitates ALPS binding (Figs. 1,2) to the same extent as conical lipids (Suppl. Fig. S2) and that conical lipids facilitate ALPS-induced membrane deformation (Fig. 3).

### Feedback mechanisms may explain the oscillating binding of ALPS and the cooperative effect of ALPS binding and conical lipids on membrane deformation

Taken together, our data suggest that membrane tension, lipid shape and membrane curvature cooperate to generate lipid packing defects and membrane deformation. Because ALPS is sensitive to membrane tension and to the presence of conical lipids and because inserting ALPS amphipathic helices increases intramembrane stresses (51), positive feedback loops could lead to the two main experimental observations we report here, osmotically-driven oscillations in ALPS binding (Fig. 4A) and DOG enhancement of ALPS-induced membrane tubulation (Fig. 4B). On a flat GUV membrane, a high tension (here driven by osmotic pressure) could facilitate ALPS binding which in turn could increase the lateral pressure in the membrane and thus further induce ALPS binding until pore formation relaxes the stress and membrane tension drops (Fig. 4A). On an initially flat membrane, conical lipids could allow ALPS binding and initiate membrane deformation through insertion of the ALPS amphipathic helix. The increase in curvature should in turn further facilitate ALPS binding and lead to membrane deformation and eventually fission (Fig. 4B).

**Figure 4.**
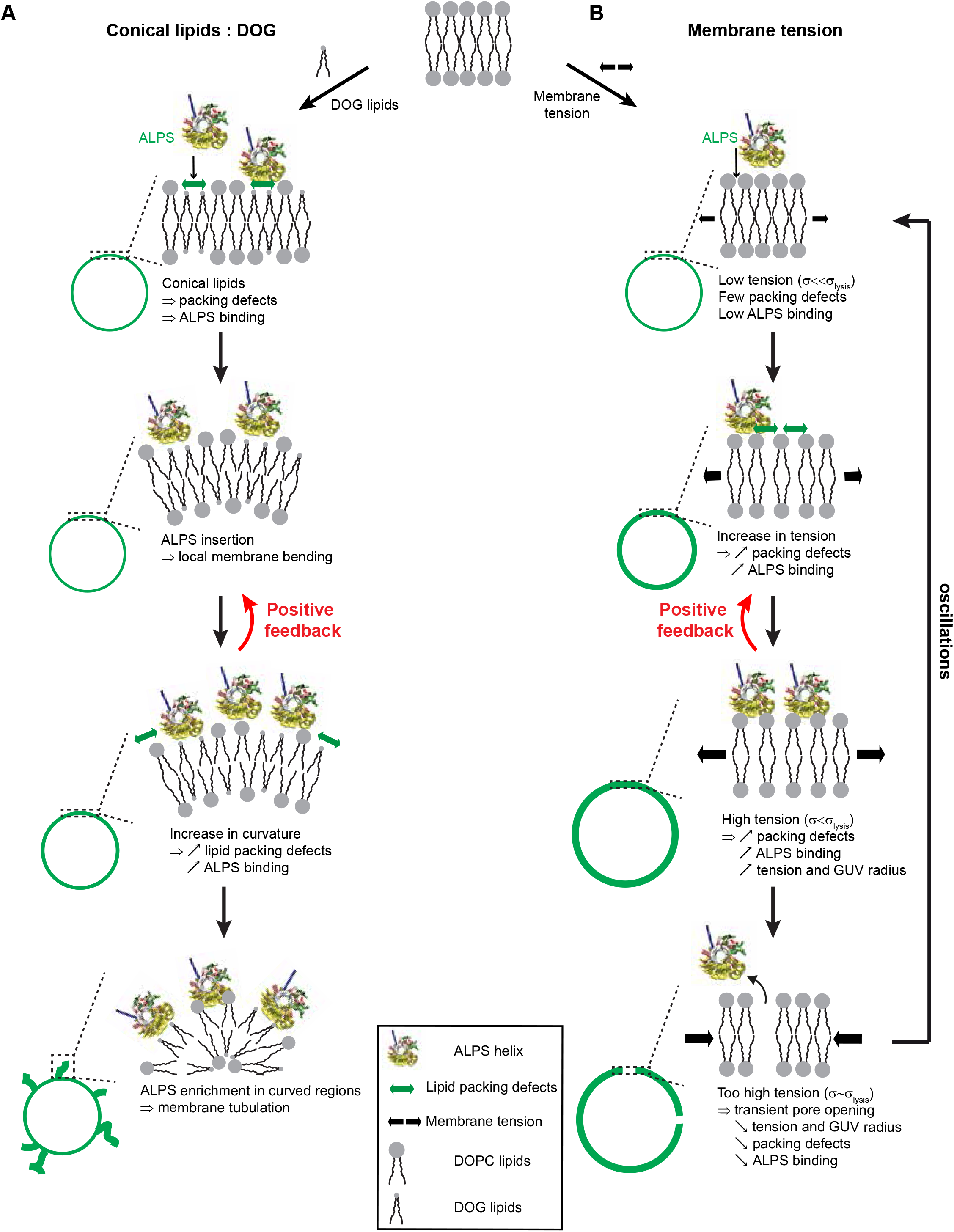
Feedback loops may explain the oscillatory binding of ALPS and the cooperative effect of ALPS binding and conical lipids on membrane deformation (A) Deformation of a flat membrane induced by positive feedback between membrane curvature and formation of lipid packing defects due to the presence of conical lipids. Conical lipids generate lipid packing defects (double green arrows) in the initially flat membrane favouring insertion of the ALPS motif in the outer leaflet of the membrane. ALPS insertion induces membrane curvature which further increases the density of lipid packing defects and ALPS binding. (B) Cyclic pore formation on a flat membrane induced by feedback between membrane tension and formation of lipid packing defects. ALPS binding increases membrane tension (black arrows) which in turn favours the formation of lipid packing defects (double green arrows) and the insertion of ALPS. When the tension reaches the critical tension for pore formation (*σ_lysis_*), a pore opens, the tension is released and ALPS detaches from the membrane.

### Tension and lipid packing

A main finding of our study is that a high membrane tension increases the density of lipid packing defects, which we show both experimentally via the binding of ALPS and through numerical simulations. Consistently, the binding to small liposomes of the N-BAR domain of *Drosophila* amphiphysin, which inserts an N-terminal amphipathic helix into the membrane, was shown to increase when membrane tension was induced osmotically (52). Numerical simulations also showed that membrane bending decreases the rigidity and the area compressibility of the liquid ordered phase in uniaxially compressed vesicles (53), suggesting that membrane tension could modify intramembrane stresses.

In both our micropipet experiments and simulations (Fig. 2), membrane tension appears to impact lipid packing above 0.1-1 mN/m, well below the lysis tension of DOPC membranes of ~10 mN/m (54). In contrast, higher tensions (3-10 mN/m) were shown to be induced by osmotic pressure differences (55), suggesting that even small differences in osmotic pressure could induce changes in lipid packing. Accordingly, we observed a strong heterogeneity in ALPS binding in a given population of DOPC GUVs. Such an heterogeneity could be due to differences in membrane tension or to the presence of DOPC oxidation products that have been shown to increase membrane permeability (56). Interestingly, oscillating phase separation was observed on GUVs which were attributed to osmotically-driven oscillations in membrane tension (57), with a feedback mechanism involving pore formation very similar to the mechanism we propose to explain oscillations in ALPS binding (Fig. 4A).

### Curvature inducers and curvature sensors

The binding of the ALPS motif of ArfGAP1 to membranes was shown previously to be extremely sensitive to lipid packing defects, hence to changes in membrane curvature. ALPS is thus thought to be an excellent ‘curvature sensor’ (19, 20, 25, 50). We show here that ALPS is not only able to sense packing defects induced by curvature or lipid shape, but also lipid packing defects induced by an increase in membrane tension. As such ALPS could also be viewed as a tension sensor. However, as ALPS inserts into the lipid bilayer, it also perturbs the lipid bilayer and generates intramembrane stresses (51, 58), that can lead to membrane deformation. We show here that ALPS can indeed be considered as a ‘curvature inducer’ (Fig. S4A-D) and that its membrane deformation activity is enhanced at high densities of lipid packing defects in the presence of conical lipids (Fig. 3D-E). Besides ALPS domain-containing proteins (20), an increasing number of membrane binding proteins have been described as ‘curvature sensors’ and/or ‘curvature inducers’, including BAR domain proteins (59) such as amphiphysin (49, 60), endophilin (61) or Arfaptin (62), proteins bearing an amphipathic helix such as epsin (63, 64), Arf1 (ADP-ribosylation factor) (65–67) and α-synuclein (68) or proteins with lipid modifications such as RAB (46) or RHO (69) family proteins, raising the question whether curvature sensing and curvature induction are always correlated. Comparing the curvature sensing properties of different proteins in a quantitative manner is possible by measuring the spontaneous curvature in lipid nanotube pulling experiments (46, 49). However, in the case of ALPS, this approach is technically challenging since ALPS does not bind to flat membranes in the absence of conical lipids or elevated tension. Comparing the ability of proteins to deform membranes is more complex since curvature induction strongly depends on protein concentration, potential multimerization states and membrane composition. Nevertheless, it is tempting to speculate that a curvature sensor will also be a curvature inducer and *vice versa.* Accordingly, the extreme sensitivity of ALPS to curvature translates into a strong ability to deform membranes (Fig. S4). In contrast, Arf1 is both a weak curvature inducer (67), a weak curvature sensor when compared to ALPS (19) and less sensitive than ALPS to lipid packing defects generated by DOG (Fig. S5 in the Supporting Material). Finally, the typical sigmoidal binding of ALPS (Fig. 3C) as previously reported for amphiphysin (49) suggests that curvature sensing dominates at low protein density while curvature induction could only occur at higher concentrations.

### Feedback loops in membrane deformation/curvature generation: relevance for intracellular trafficking

Our data suggest that membrane tension and conical lipids amplify protein binding by positive feedback on lipid packing defects, which could induce membrane deformation and eventually fission. Intracellular membranes can be separated in two main membrane ‘territories’, the early secretory pathway comprising the nuclear envelope, the endoplasmic reticulum (ER) and the cis-Golgi, and the late secretory pathway comprising the *trans-* Golgi, endosomal membranes and the plasma membrane (70–72). The first territory is characterized by loose packing and low surface charge; amphipathic helices inserts into membranes mostly via hydrophobic interactions. The second territory displays a low density of packing defects and a high surface charges; helix insertion occurs through electrostatic interactions. Conical lipids, by generating loose lipid packing, could selectively target ALPS domain proteins to membranes in the early secretory pathway. Accordingly, at the cis-Golgi, the golgin GMAP-210 selects vesicles with loose lipid packing through the binding of its ALPS motif (73). Production of DAGs by phospholipases could locally increase the density of lipid packing defects and, consequently, the local concentration of ALPS domain proteins could reach their concentration threshold for membrane deformation. Alternative mechanisms that could increase local protein concentration and lead to membrane bending include lipid clustering (60) and protein crowding (74).

While a high tension could help recruiting membrane bending proteins at initial stages of membrane deformation, it should inhibit bud formation by coat proteins at later stages (67, 75). The role of tension in scission is also complex. Intuitively, tension should facilitate membrane fission(76). Accordingly, pulling forces trigger the scission of membrane tubes scaffolded by BAR proteins through a friction-driven mechanism in dynamin-independent fission (77). However, in dividing cells, membrane tension drops before the final abscission step of cytokinesis (78). While membrane tension at the cell plasma membrane has been extensively studied in different biological contexts and in particular cell motility and adhesion (79, 80), membrane tension of intracellular compartments remains largely elusive. *In vitro* reconstitution of ER and Golgi membrane networks shows that the tension of ER tubules is higher than that of Golgi tubules (81). A gradient of membrane tension may exist along the secretory pathway which could generate higher densities of lipid packing defects in the early secretory pathway and favour the binding of proteins sensitive to lipid packing such as ALPS domain proteins to ER or cis-Golgi membranes.

## Supporting material

Supporting materials and methods, five figures and five movies are available.

## Author contributions

M.P., S.V., B.G. and J.-B.M. designed the experiments; M.P., E.A., S.V. and D.G. performed the experiments and analysis; M.P. and J.-B.M. wrote the article, with assistance from the co-authors.

## Acknowledgements

We thank Patricia Bassereau, Sophie Aimon and Gil Toombes for assistance with image processing, Bruno Antonny and Guillaume Drin for discussions and help with ALPS purification. S.V. acknowledges support by the Swiss National Science Foundation (#163966). M.P was funded by a grant from the French Ministry of Research. Other fundings are from Institut Curie, CNRS (Program ‘Prise de risque à l’interface Physique-Biologie’), ANR (grant number ANR09-JCJC-0020-01), INSERM Plan Cancer 2009-2013 INSERM - CEA Tecsan (grant number PC201125). E.A. is a Career Member of CONICET (Argentina). E.A. thanks CONICET for the financial support (PIP 2013-2015 grant). The authors declare no conflict of interest.

## References

1. Turing, A.M. 1952. The Chemical Basis of Morphogenesis. Philos. Trans. R. Soc. London B Biol. Sci. 237.

2. Loose, M., E. Fischer-Friedrich, J. Ries, K. Kruse, and P. Schwille. 2008. Spatial Regulators for Bacterial Cell Division Self-Organize into Surface Waves in Vitro. Science (80-.). 320: 789–792.

3. Vecchiarelli, A.G., M. Li, M. Mizuuchi, L.C. Hwang, Y. Seol, K.C. Neuman, and K. Mizuuchi. 2016. Membrane-bound MinDE complex acts as a toggle switch that drives Min oscillation coupled to cytoplasmic depletion of MinD. Proc. Natl. Acad. Sci. U. S. A. 113: E1479–88.

4. Bonazzi, D., A. Haupt, H. Tanimoto, D. Delacour, D. Salort, and N. Minc. 2015. Actin-Based Transport Adapts Polarity Domain Size to Local Cellular Curvature. Curr. Biol. 25: 2677–2683.

5. Martin, S.G. 2015. Spontaneous cell polarization: Feedback control of Cdc42 GTPase breaks cellular symmetry. BioEssays. 37: 1193–1201.

6. Terenna, C.R., T. Makushok, G. Velve-Casquillas, D. Baigl, Y. Chen, M. Bornens, A. Paoletti, M. Piel, and P.T. Tran. 2008. Physical Mechanisms Redirecting Cell Polarity and Cell Shape in Fission Yeast. Curr. Biol. 18: 1748–1753.

7. Chaix, A., A. Zarrinpar, and S. Panda. 2016. The circadian coordination of cell biology. J. Cell Biol. 215: 15–25.

8. Yang, Q., and J.E. Ferrell. 2013. The Cdk1–APC/C cell cycle oscillator circuit functions as a time-delayed, ultrasensitive switch. Nat. Cell Biol. 15: 519–525.

9. Jackson, C.L. 2014. GEF-effector interactions. Cell. Logist. 4: e943616.

10. Kozlov, M.M., F. Campelo, N. Liska, L. V Chernomordik, S.J. Marrink, and H.T. McMahon. 2014. Mechanisms shaping cell membranes. Curr. Opin. Cell Biol. 29: 53–60.

11. Pinot, M., B. Goud, and J.-B. Manneville. 2010. Physical aspects of COPI vesicle formation. Mol. Membr. Biol. 27: 428–42.

12. Bigay, J., and B. Antonny. 2012. Curvature, Lipid Packing, and Electrostatics of Membrane Organelles: Defining Cellular Territories in Determining Specificity. Dev. Cell. 23: 886–895.

13. Liu, J., Y. Sun, D.G. Drubin, and G.F. Oster. 2009. The Mechanochemistry of Endocytosis. PLoS Biol. 7: e1000204.

14. Taylor, M.J., M. Lampe, and C.J. Merrifield. 2012. A Feedback Loop between Dynamin and Actin Recruitment during Clathrin-Mediated Endocytosis. PLoS Biol. 10: e1001302.

15. Wang, X., B.J. Galletta, J.A. Cooper, and A.E. Carlsson. 2016. Actin-Regulator Feedback Interactions during Endocytosis. Biophys. J. 110: 1430–1443.

16. Lieber, A.D., S. Yehudai-Resheff, E.L. Barnhart, J.A. Theriot, and K. Keren. 2013. Membrane Tension in Rapidly Moving Cells Is Determined by Cytoskeletal Forces. Curr. Biol. 23: 1409–1417.

17. Tsujita, K., T. Takenawa, and T. Itoh. 2015. Feedback regulation between plasma membrane tension and membrane-bending proteins organizes cell polarity during leading edge formation. Nat. Cell Biol. 17: 749–758.

18. Bigay, J., J.F. Casella, G. Drin, B. Mesmin, and B. Antonny. 2005. ArfGAP1 responds to membrane curvature through the folding of a lipid packing sensor motif. EMBO J. 24: 2244–2253.

19. Ambroggio, E., B. Sorre, P. Bassereau, B. Goud, J.-B.B. Manneville, and B. Antonny. 2010. ArfGAP1 generates an Arf1 gradient on continuous lipid membranes displaying flat and curved regions. EMBO J. 29: 292–303.

20. Antonny, B. 2011. Mechanisms of membrane curvature sensing. Annu. Rev. Biochem. 80: 101–23.

21. Cui, H., E. Lyman, and G. a Voth. 2011. Mechanism of membrane curvature sensing by amphipathic helix containing proteins. Biophys. J. 100: 1271–9.

22. Vamparys, L., R. Gautier, S. Vanni, W.F.D. Bennett, D.P. Tieleman, B. Antonny, C. Etchebest, and P.F.J. Fuchs. 2013. Conical Lipids in Flat Bilayers Induce Packing Defects Similar to that Induced by Positive Curvature. Biophys. J. 104: 585–593.

23. Vanni, S., H. Hirose, H. Barelli, B. Antonny, and R. Gautier. 2014. A sub-nanometre view of how membrane curvature and composition modulate lipid packing and protein recruitment. Nat. Commun. 5: 4916.

24. Risselada, H.J., and S.J. Marrink. 2009. Curvature effects on lipid packing and dynamics in liposomes revealed by coarse grained molecular dynamics simulations. Phys. Chem. Chem. Phys. 11: 2056.

25. Vanni, S., L. Vamparys, R. Gautier, G. Drin, C. Etchebest, P.F.J. Fuchs, and B. Antonny. 2013. Amphipathic Lipid Packing Sensor Motifs: Probing Bilayer Defects with Hydrophobic Residues. Biophys. J. 104: 575–584.

26. Angelova, M.I., S. Soléau, P. Méléard, J.F. Faucon, and P. Bothorel. 1992. Preparation of giant vesicles by external AC electric fields. Kinetics and applications. 89: 127–131.

27. Manneville, J.-B., C. Leduc, B. Sorre, and G. Drin. 2012. Studying In Vitro Membrane Curvature Recognition by Proteins and its Role in Vesicular Trafficking. Methods Cell Biol.

28. Evans, E., and W. Rawicz. 1990. Entropy-driven tension and bending elasticity in condensed-fluid membranes. Phys Rev Lett. 64: 2094–2097.

29. Shinoda, W., R. DeVane, and M.L. Klein. 2007. Multi-property fitting and parameterization of a coarse grained model for aqueous surfactants. Mol. Simul. 33: 27–36.

30. Shinoda, W., R. DeVane, and M.L. Klein. 2010. Zwitterionic lipid assemblies: molecular dynamics studies of monolayers, bilayers, and vesicles using a new coarse grain force field. J. Phys. Chem. B. 114: 6836–49.

31. MacDermaid, C.M., H.K. Kashyap, R.H. DeVane, W. Shinoda, J.B. Klauda, M.L. Klein, and G. Fiorin. 2015. Molecular dynamics simulations of cholesterol-rich membranes using a coarse-grained force field for cyclic alkanes. J. Chem. Phys. 143: 243144.

32. Plimpton, S. 1995. Fast Parallel Algorithms for Short-Range Molecular Dynamics. J. Comput. Phys. 117: 1–19.

33. Nosé, S. 1984. A molecular dynamics method for simulations in the canonical ensemble. Mol. Phys. 52: 255–268.

34. Parrinello, M., and A. Rahman. 1981. Polymorphic transitions in single crystals: A new molecular dynamics method. J. Appl. Phys. 52: 7182–7190.

35. Martyna, G.J., D.J. Tobias, and M.L. Klein. 1994. Constant pressure molecular dynamics algorithms. J. Chem. Phys. 101: 4177–4189.

36. Eastwood, J.W., R.W. Hockney, and D.N. Lawrence. 1980. P3M3DP—The three-dimensional periodic particle-particle/ particle-mesh program. Comput. Phys. Commun. 19: 215–261.

37. Humphrey, W., A. Dalke, and K. Schulten. 1996. VMD: Visual molecular dynamics. J. Mol. Graph. 14: 33–38.

38. Irving, J.H., and J.G. Kirkwood. 1950. The Statistical Mechanical Theory of Transport Processes. IV. The Equations of Hydrodynamics. J. Chem. Phys. 18: 817–829.

39. Safouane, M., L. Berland, A. Callan-Jones, B. Sorre, W. Römer, L. Johannes, G.E.S. Toombes, P. Bassereau, and et al. 2010. Lipid co-sorting mediated by Shiga toxin induced tubulation. Traffic. 11: 1519–29.

40. Antonny, B., I. Huber, S. Paris, M. Chabre, and D. Cassel. 1997. Activation of ADP-ribosylation factor 1 GTPase-activating protein by phosphatidylcholine-derived diacylglycerols. J. Biol. Chem. 272: 30848–51.

41. Sandre, O., L. Moreaux, and F. Brochard-Wyart. 1999. Dynamics of transient pores in stretched vesicles. Proc. Natl. Acad. Sci. U. S. A. 96: 10591–6.

42. Karatekin, E., O. Sandre, H. Guitouni, N. Borghi, P.-H. Puech, and F. Brochard-Wyart. 2003. Cascades of transient pores in giant vesicles: line tension and transport. Biophys. J. 84: 1734–49.

43. Oglęcka, K., P. Rangamani, B. Liedberg, R.S. Kraut, and A.N. Parikh. 2014. Oscillatory phase separation in giant lipid vesicles induced by transmembrane osmotic differentials. Elife. 3: e03695.

44. Bacle, A., R. Gautier, C.L. Jackson, P.F.J. Fuchs, and S. Vanni. 2017. Interdigitation between Triglycerides and Lipids Modulates Surface Properties of Lipid Droplets. Biophys. J. 112: 1417–1430.

45. Pinot, M., S. Vanni, S. Pagnotta, S. Lacas-Gervais, L.-A. Payet, T. Ferreira, R. Gautier, B. Goud, B. Antonny, and H. Barelli. 2014. Lipid cell biology. Polyunsaturated phospholipids facilitate membrane deformation and fission by endocytic proteins. Science. 345: 693–7.

46. Kulakowski, G., H. Bousquet, J.-B. Manneville, P. Bassereau, B. Goud, and L.K. Oesterlin. 2018. Lipid packing defects and membrane charge control RAB GTPase recruitment. Traffic.

47. Holopainen, J.M., M.I. Angelova, T. Söderlund, and P.K.J. Kinnunen. 2002. Macroscopic consequences of the action of phospholipase C on giant unilamellar liposomes. Biophys. J. 83: 932–43.

48. Sheetz, M.P., and S.J. Singer. 1974. Biological membranes as bilayer couples. A molecular mechanism of drug-erythrocyte interactions. Proc Natl Acad Sci U S A. 71: 4457–4461.

49. Sorre, B., A. Callan-Jones, J. Manzi, B. Goud, J. Prost, P. Bassereau, and A. Roux. 2011. Nature of curvature coupling of amphiphysin with membranes depends on its bound density. Proc. Natl. Acad. Sci. U. S. A. 109: 173–8.

50. Bigay, J., P. Gounon, S. Robineau, B. Antonny, D. Deangelis, and J. Rothman. 2003. Lipid packing sensed by ArfGAP1 couples COPI coat disassembly to membrane bilayer curvature. Nature. 426: 563–566.

51. Campelo, F., and M.M. Kozlov. 2014. Sensing Membrane Stresses by Protein Insertions. PLoS Comput. Biol. 10: e1003556.

52. Hutchison, J.B., A.P.K.K. Karunanayake Mudiyanselage, R.M. Weis, and A.D. Dinsmore. 2016. Osmotically-induced tension and the binding of N-BAR protein to lipid vesicles. Soft Matter. 12: 2465–2472.

53. Risselada, H., S. Marrink, and M. Müller. 2011. Curvature-Dependent Elastic Properties of Liquid-Ordered Domains Result in Inverted Domain Sorting on Uniaxially Compressed Vesicles. Phys. Rev. Lett. 106: 8–11.

54. Olbrich, K., W. Rawicz, D. Needham, and E. Evans. 2000. Water permeability and mechanical strength of polyunsaturated lipid bilayers. Biophys. J. 79: 321–7.

55. Alam Shibly, S.U., C. Ghatak, M.A. Sayem Karal, M. Moniruzzaman, and M. Yamazaki. 2016. Experimental Estimation of Membrane Tension Induced by Osmotic Pressure. Biophys. J. 111: 2190–2201.

56. Heuvingh, J., and S. Bonneau. 2009. Asymmetric oxidation of giant vesicles triggers curvature-associated shape transition and permeabilization. Biophys. J. 97: 2904–12.

57. Oglęcka, K., P. Rangamani, B. Liedberg, R.S. Kraut, and A.N. Parikh. 2014. Oscillatory phase separation in giant lipid vesicles induced by transmembrane osmotic differentials. Elife. 3: e03695.

58. Campelo, F., H.T. McMahon, and M.M. Kozlov. 2008. The hydrophobic insertion mechanism of membrane curvature generation by proteins. Biophys. J. 95: 2325–39.

59. Simunovic, M., G.A. Voth, A. Callan-Jones, and P. Bassereau. 2015. When Physics Takes Over: BAR Proteins and Membrane Curvature. Trends Cell Biol. 25: 780–792.

60. Picas, L., J. Viaud, K. Schauer, S. Vanni, K. Hnia, V. Fraisier, A. Roux, P. Bassereau, F. Gaits-Iacovoni, B. Payrastre, J. Laporte, J.-B. Manneville, and B. Goud. 2014. BIN1/M-Amphiphysin2 induces clustering of phosphoinositides to recruit its downstream partner dynamin. Nat. Commun. 5.

61. Simunovic, M., E. Evergren, I. Golushko, C. Prévost, H.-F. Renard, L. Johannes, H.T. McMahon, V. Lorman, G.A. Voth, and P. Bassereau. 2016. How curvature-generating proteins build scaffolds on membrane nanotubes. Proc. Natl. Acad. Sci. 113: 11226–11231.

62. Ambroggio, E.E., J. Sillibourne, B. Antonny, J.-B. Manneville, and B. Goud. 2013. Arf1 and Membrane Curvature Cooperate to Recruit Arfaptin2 to Liposomes. PLoS One. 8: e62963.

63. Ford, M.G.J., I.G. Mills, B.J. Peter, Y. Vallis, G.J.K. Praefcke, P.R. Evans, and H.T. McMahon. 2002. Curvature of clathrin-coated pits driven by epsin. Nature. 419: 361–6.

64. Capraro, B.R., Y. Yoon, W. Cho, and T. Baumgart. 2010. Curvature sensing by the epsin N-terminal homology domain measured on cylindrical lipid membrane tethers. J. Am. Chem. Soc. 132: 1200–1.

65. Krauss, M., J.-Y. Jia, A. Roux, R. Beck, F.T. Wieland, P. De Camilli, and V. Haucke. 2008. Arf1-GTP-induced tubule formation suggests a function of Arf family proteins in curvature acquisition at sites of vesicle budding. J. Biol. Chem. 283: 27717–23.

66. Beck, R., Z. Sun, F. Adolf, C. Rutz, J. Bassler, K. Wild, I. Sinning, E. Hurt, B. Brugger, J. Bethune, and F. Wieland. 2008. Membrane curvature induced by Arf1-GTP is essential for vesicle formation. Proc Natl Acad Sci U S A. 105: 11731–11736.

67. Manneville, J.-B., J.-F. Casella, E. Ambroggio, P. Gounon, J. Bertherat, P. Bassereau, J. Cartaud, B. Antonny, and B. Goud. 2008. COPI coat assembly occurs on liquid-disordered domains and the associated membrane deformations are limited by membrane tension. Proc. Natl. Acad. Sci. U. S. A. 105: 16946–51.

68. Pranke, I.M., V. Morello, J. Bigay, K. Gibson, J.-M. Verbavatz, B. Antonny, and C.L. Jackson. 2011. {alpha}-Synuclein and ALPS motifs are membrane curvature sensors whose contrasting chemistry mediates selective vesicle binding. J. Cell Biol. 194: 89–103.

69. Park, S.-Y., J.-S. Yang, A.B. Schmider, R.J. Soberman, and V.W. Hsu. 2015. Coordinated regulation of bidirectional COPI transport at the Golgi by CDC42. Nature. 521: 529–532.

70. Bigay, J., and B. Antonny. 2012. Curvature, Lipid Packing, and Electrostatics of Membrane Organelles: Defining Cellular Territories in Determining Specificity. Dev. Cell. 23: 886–895.

71. Holthuis, J.C.M., and A.K. Menon. 2014. Lipid landscapes and pipelines in membrane homeostasis. Nature. 510: 48–57.

72. Jackson, C.L., L. Walch, and J.-M. Verbavatz. 2016. Lipids and Their Trafficking: An Integral Part of Cellular Organization. Dev. Cell. 39: 139–153.

73. Magdeleine, M., R. Gautier, P. Gounon, H. Barelli, S. Vanni, and B. Antonny. 2016. A filter at the entrance of the Golgi that selects vesicles according to size and bulk lipid composition. EMBO J. 5: 292–303.

74. Stachowiak, J.C., E.M. Schmid, C.J. Ryan, H.S. Ann, D.Y. Sasaki, M.B. Sherman, P.L. Geissler, D.A. Fletcher, and C.C. Hayden. 2012. Membrane bending by protein–protein crowding. Nat. Cell Biol. 14: 944–949.

75. Saleem, M., S. Morlot, A. Hohendahl, J. Manzi, M. Lenz, and A. Roux. 2015. A balance between membrane elasticity and polymerization energy sets the shape of spherical clathrin coats. Nat. Commun. 6: 6249.

76. Roux, A., K. Uyhazi, A. Frost, and P. De Camilli. 2006. GTP-dependent twisting of dynamin implicates constriction and tension in membrane fission. Nature. 441: 528–531.

77. Simunovic, M., J.-B. Manneville, H.-F. Renard, E. Evergren, K. Raghunathan, D. Bhatia, A.K. Kenworthy, G.A. Voth, J. Prost, H.T. McMahon, L. Johannes, P. Bassereau, and A. Callan-Jones. 2017. Friction Mediates Scission of Tubular Membranes Scaffolded by BAR Proteins. Cell. 170.

78. Lafaurie-Janvore, J., P. Maiuri, I. Wang, M. Pinot, J.-B. Manneville, T. Betz, M. Balland, and M. Piel. 2013. ESCRT-III assembly and cytokinetic abscission are induced by tension release in the intercellular bridge. Science (80-. ). 340.

79. Gauthier, N.C., T.A. Masters, and M.P. Sheetz. 2012. Mechanical feedback between membrane tension and dynamics. Trends Cell Biol. 22: 527–35.

80. Keren, K. 2011. Cell motility: the integrating role of the plasma membrane. Eur. Biophys. J.: 1013–1027.

81. Upadhyaya, A., and M.P. Sheetz. 2004. Tension in tubulovesicular networks of Golgi and endoplasmic reticulum membranes. Biophys J. 86: 2923–2928.

